# Using an Optimal Set of Features with a Machine Learning-Based Approach to Predict Effector Proteins for *Legionella pneumophila*

**DOI:** 10.1101/383570

**Authors:** Zhila Esna Ashari, Kelly A. Brayton, Shira L. Broschat

**Affiliations:** School of Electrical Engineering and Computer Science, Washington State University, Pullman, Washington, United States; Department of Veterinary Microbiology and Pathology, Washington State University, Washington, United States; Paul G. Allen School for Global Animal Health, Washington State University, Pullman, Washington, United States

## Abstract

Type IV secretion systems exist in a number of bacterial pathogens and are used to secrete effector proteins directly into host cells in order to change their environment making the environment hospitable for the bacteria. In recent years, several machine learning algorithms have been developed to predict effector proteins, potentially facilitating experimental verification. However, inconsistencies exist between their results. Previously we analysed the disparate sets of predictive features used in these algorithms to determine an optimal set of 370 features for effector prediction. This work focuses on the best way to use these optimal features by designing three machine learning classifiers, comparing our results with those of others, and obtaining de novo results. We chose the pathogen *Legionella pneumophila* strain Philadelphia-1, a cause of Legionnaires’ disease, because it has many validated effector proteins and others have developed machine learning prediction tools for it. While all of our models give good results indicating that our optimal features are quite robust, Model 1, which uses all 370 features with a support vector machine, has slightly better accuracy. Moreover, Model 1 predicted 760 effector proteins, more than any other study, 315 of which have been validated. Although the results of our three models agree well with those of other researchers, their models only predicted 126 and 311 candidate effectors.

## Introduction

Bacterial pathogens can use secretion systems to deliver proteins to the host cell. There are nine known secretion systems, but the focus of this work is on the type IV secretion system (T4SS). The T4SS is composed of multiple proteins responsible for secreting effector proteins directly into eukaryotic host cells. When effector proteins are translocated into host cells, they manipulate their defence systems, causing infections. In order to understand how these effector proteins manipulate the host cell, it is first necessary to identify them. However, this can be a difficult task because they are not well conserved among organisms. Several methods have been proposed for identifying effector proteins with experimental validation being the most accurate but also the most expensive and time consuming [1-4]. Accurate prediction of candidate effectors would expedite the experimental validation process. As a result, recent studies have focused on using prediction approaches such as scoring effector proteins based on their characteristics or using machine learning algorithms [5-10]. Because these methods considered different sets of features, we examined their effectiveness in an earlier study and determined a set of optimal features for prediction of T4SS effector proteins [11-12]. By features, we refer here to the characteristics and properties of protein sequences that can be measured and thus assigned binary or continuous numerical values.

In our previous study, we identified a set of optimal features using four datasets of validated effector and non-effector proteins from four different Proteobacterial pathogens, *Legionella pneumophila, Coxiella burnettii, Bartonella* spp., and *Brucella* spp. that works well for prediction of T4SS effector proteins. In this work, we use this set of optimal features to develop a machine learning based classifier to predict T4SS effectors, which is trained using the set of validated effector and non-effector proteins from our earlier study of all four pathogens. Our goals are four-fold: i) to test our classifier on a pathogen with many validated effectors to ascertain how well it works for a single pathogen, ii) to determine the best way to use the optimal features to achieve the most accurate results, iii) to compare our results with those of other T4SS effector prediction models, and iv) to obtain de novo results. Therefore, we selected the *L. pneumophila* strain Philadelphia-1 genome/deduced proteome as the subject of our study because it has the greatest number of validated effector proteins, and several prediction algorithms have used this organism as their subject. *L. pneumophila* is a Gram-negative bacterial pathogen from the class Gammaproteobacteria which causes Legionnaires’ disease, and many researches have focused on this pathogen and its effector proteins [13-27].

To analyse our optimal features, we actually developed three different machine learning classifiers. We first explain how we design and validate our three machine learning models, two of which are ensemble classifiers. Next, we use the models on the whole proteome from *L. pneumophila* strain Philadelphia-1 and compare our results with those of previous studies for *L. pneumophila.* Finally, we obtain de novo predictions of effector proteins for *L. pneumophila.*

## Materials and Methods

Fig. 1 represents the workflow used to complete this study. Each step is described in more detail in subsequent sections.

**Figure 1.**
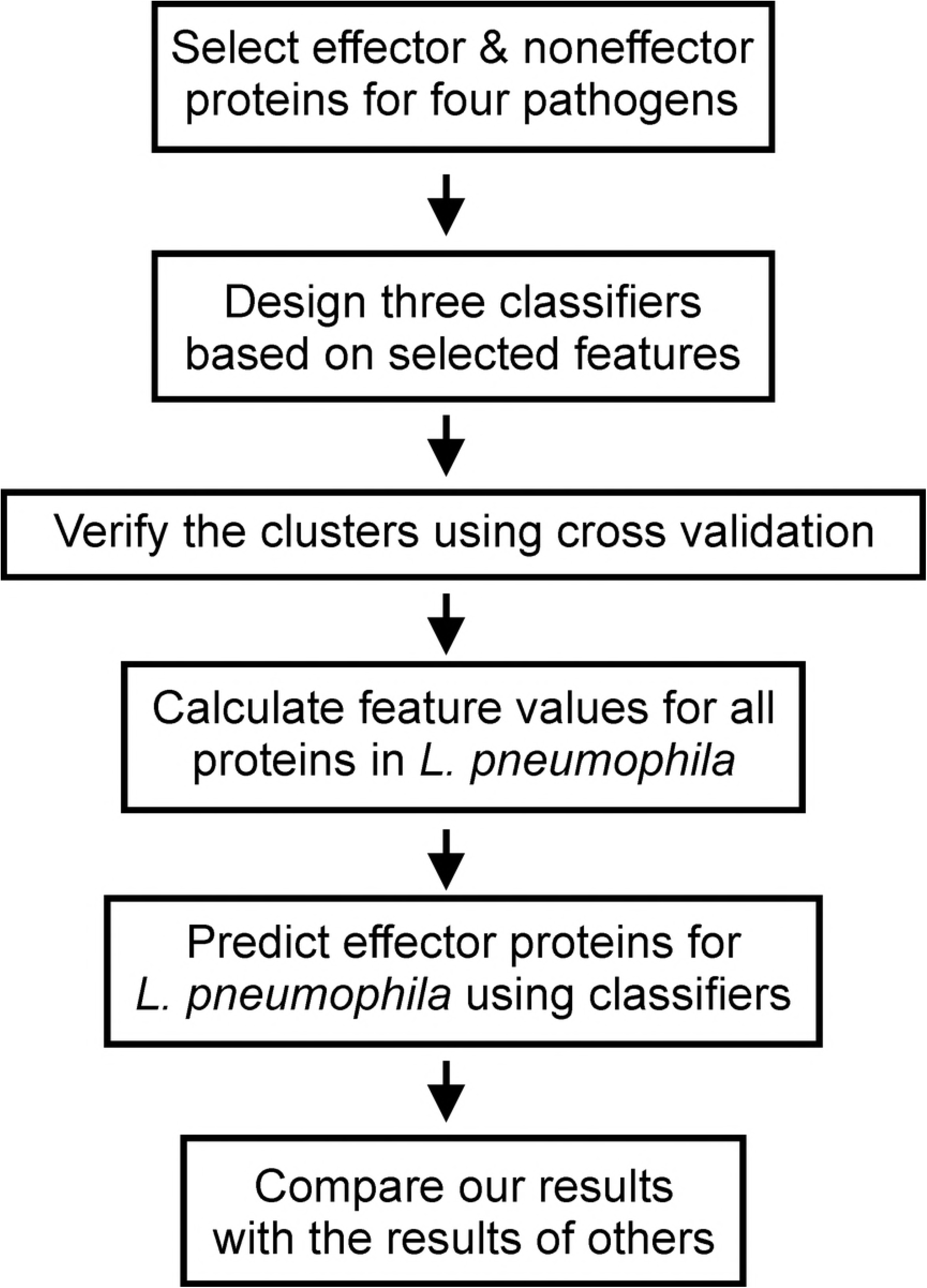
Workflow.

### (a) Creating training and test datasets

Our training dataset was composed of effectors and non-effectors from four different bacterial pathogens: *L. pneumophila, C. burnettii, Brucella* spp., and *Bartonella* spp. In our previous paper, each of these pathogens was treated as a separate dataset [12], and we determined effective features for each using a feature selection method. Based on our results, we proposed a final set of effective features for prediction of T4SS effectors. In the present study we merged these four datasets to create a set of known effectors and non-effectors which was used as the training set for our problem. This dataset consisted of 1,127 data points among which there were 429 effectors and 698 non-effectors. The protein sequences for our training dataset are presented in [S1 File]. Moreover, we created a test set, which is composed of 2,942 protein sequences from the complete proteome of *L. pneumophila* strain Philadelphia-1 [S2 File].

### (b) Features

The features used in this work are the set of optimal features proposed in our earlier work [12]. In our previous study we did a comprehensive literature review and compiled a list of all the features used for prediction of T4SS effector proteins. Because some of the features were vectors, we began with 1,027 features. By vector, we mean that a particular feature had multiple values. For example, there are 20 different amino acids so that the amino acid composition feature for a protein sequence has 20 different percentage values. Using a multi-level feature selection approach, we proposed a set of optimal features for our prediction problem and retained 370 features. Overall, they include chemical properties, structural properties, compositional properties, and position-specific scoring matrix (PSSM)-related properties, which are a type of compositional property.

Our optimal feature set includes 15 features that are related to the chemical and structural properties of protein sequences. Chemical properties such as hydropathy are considered to be important for T4SS effector prediction because they determine how proteins interact with their environment and because they are believed to be key mediators in determining how effectors enter host cells [6, 8]. The structural properties of proteins, such as coiled coil domains, allow protein-protein interactions within host cells thus effecting cellular processes [6, 8-9]. Our feature set also includes compositional properties of protein sequences, comprising selected elements of the amino acid and dipeptide composition vectors totalling 57 in number. In addition, they include 298 features from the PSSM profile for protein sequences and its auto-covariance correlation composition vector [28]. Compositional properties are considered to be effective for T4SS effector prediction because they determine the shape of the protein, and they also account for amino acid frequencies and motifs [7].

All features are explained at greater length in [11].

### (c) Machine Learning Models and Validation

A major goal of this paper was to determine how to use the optimal feature set to obtain the most accurate results. As such, we considered different methodologies and algorithms, for example, using a single classifier versus an ensemble classifier, and decided to design three separate models based on a division of the features. To test our classifiers, we used several standard metrics for machine learning models: accuracy, recall, and precision.

Our first model, Model 1, was based on the use of the entire optimal feature set. We calculated the features for all the protein sequences in our dataset of effectors and non-effectors. These 370 features are shown in [S1 Table]. We used this dataset to train a support vector machine (SVM) classifier. An SVM is a powerful machine learning classifier often used for supervised learning, that is learning based on using labelled training data [29]. It allows the use of different Kernel functions to create classifiers that fit a dataset. Our second and third models, Models 2 and 3, were ensemble classifiers composed of three separate classifiers. Each of these classifiers was designed to work with a subset of the optimal feature set. By dividing the features among several classifiers, we wanted to decrease the possibility of overfitting effects on our results. Overfitting occurs when a model fits training data too well, causing the model to be less accurate for new data. Here, we chose three SVM classifiers for each ensemble model and with all redundant and highly correlated features removed; each of three SVM classifiers determines whether a protein sequence was an effector protein or a non-effector protein. The final prediction was based on the output class that had the majority of votes from all three classifiers. When two or more classifiers voted for a protein sequence to be an effector, it was predicted to be an effector protein. We used the SVM tuning function in R to find the best parameters for our SVM classifiers which resulted in the use of a radial Kernel and a C parameter of 1 [30].

As mentioned, Model 1 used all the selected features. For our first ensemble classifier, Model 2, the three groups of features were divided among our three classifiers as follows: i) features related to PSSM composition, ii) features related to the auto-covariance correlation of PSSM, and iii) chemical, structural, and compositional features [S1 Table] (e.g., amino acid composition, dipeptide composition, average hydropathy, total hydropathy, hydropathy of C terminal, hydropathy of N terminal, number of coiled coil regions, signal peptide probability, polarity, molecular mass, length, and homology to known effectors). For our second ensemble classifier, Model 3, the three groups of features divided among our classifiers were as follows: i) PSSM-related features (PSSM composition and auto covariance correlation of PSSM), ii) features related to the composition of amino acids in protein sequences (amino acid composition and dipeptide composition), and iii) chemical and structural features (average hydropathy, total hydropathy, hydropathy of C terminal, hydropathy of N terminal, number of coiled coil regions, signal peptide probability, polarity, molecular mass, length, and homology to known effectors).

After building our dataset and designing our machine learning classifiers, we used 10-fold cross-validation to validate our models and to test for overfitting in the results. The dataset was randomly divided into ten groups, and for each fold, one group was kept for testing and the other nine groups were used for training. We calculated confusion matrices for each cross-validation step for all three models. A confusion matrix is a table that displays the results of a machine learning algorithm for known test data. When a positive value (here an effector protein) is correctly identified, it is called a true positive (TP); when a negative value (here a non-effector protein) is correctly identified, it is called a true negative (TN); when a positive value is identified as a negative value, it is called a false negative (FN); and when a negative value is identified as a positive value, it is called a false positive (FP). From the confusion matrices, we calculated accuracy measures for the models. The final accuracy for the models was obtained by taking the average of the ten different folds. In addition, because the number of effectors (429) and non-effectors (698) in our dataset was not the same, we calculated recall and precision. Recall is a measure of sensitivity, and precision is a measure of relevance. When these values are sufficiently high, it indicates that our results are not affected by the unbalanced dataset and are another indication of the accuracy of the results. The equations for accuracy, recall, and precision are presented in (1)–(3) [31].

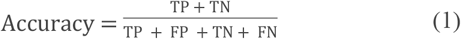

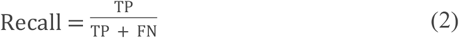

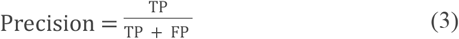

The next step after designing and validating our models was to use them for prediction of effector proteins in the whole proteome of *L. pneumophila* strain Philadelphia-1. This proteome contains 2,942 protein sequences and was used as our test set [S2 File]. We calculated the feature values for all the protein sequences in *L. pneumophila* using different tools and programming languages as described in [11]. We then used our three models for de novo prediction of effector proteins in the *L. pneumophila* proteome. Models 2 and 3 each consisted of 3 separate classifiers with each classifier determining whether one of the 2,942 *L. pneumophila* protein sequences was an effector or non-effector. Protein sequences receiving two or three positive votes were predicted as effectors.

The final step in this study was to compare our results to those obtained previously by others for prediction of effector proteins for *L. pneumophila.* We selected the study performed by Burstein et al. in 2009 which used a voting scheme based on four different algorithms [5] and the study performed by Meyer et al. in 2013 which used a scoring method [6]. Results and comparisons are discussed in the next section.

## Results and Discussion

We developed three models to test the accuracy of our optimal feature set. Model 1 used the entire set of 370 features with an SVM, and Models 2 and 3 also used the entire set of features. However, they were divided into subsets and used with three separate SVM classifiers comprising ensemble models. We used 10-fold cross-validation to test these models. The accuracy results calculated for each of the 10 folds are shown in Tables 1 through 3 for Models 1 through 3, respectively.

**Table 1.** Accuracy measures for 10-fold cross-validation of Model 1 using the entire feature set for prediction

|  | Fold Accuracy (%) |  |  |  |  |  |  |  |  |  |
| --- | --- | --- | --- | --- | --- | --- | --- | --- | --- | --- |
|  | 1 | 2 | 3 | 4 | 5 | 6 | 7 | 8 | 9 | 10 |
| **Model 1** | 95.13 | 93.80 | 93.75 | 92.47 | 93.75 | 93.36 | 95.08 | 95.13 | 95.11 | 92.92 |

**Table 2.**
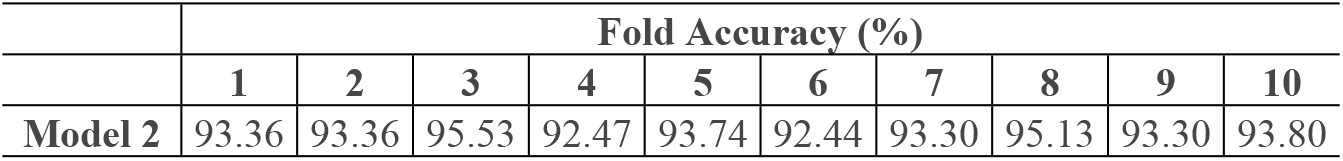
Accuracy measures for 10-fold cross-validation of Model 2 using three feature subsets: i) PSSM composition features, ii) PSSM auto-covariance correlation features, and iii) chemical, structural, and compositional features

**Table 3.** Accuracy measures for 10-fold cross-validation of Model 3 using three feature subsets: i) PSSM-related features, ii) compositional features, and iii) chemical and structural features

|  | Fold Accuracy (%) |  |  |  |  |  |  |  |  |  |
| --- | --- | --- | --- | --- | --- | --- | --- | --- | --- | --- |
|  | 1 | 2 | 3 | 4 | 5 | 6 | 7 | 8 | 9 | 10 |
| **Model 3** | 90.70 | 91.59 | 92.41 | 91.59 | 94.64 | 92.92 | 93.30 | 90.13 | 93.30 | 93.80 |

**Table 4.** Average accuracy, recall, and precision measures over 10 folds for the three effector prediction models

|  | **Model 1** | **Model 2** | **Model 3** |
| --- | --- | --- | --- |
| **Average accuracy** | 94.05% | 93.64% | 92.44% |
| **Average recall** | 92.00% | 93.06% | 92.83% |
| **Average precision** | 92.49% | 90.91% | 87.33% |

The three values are 94.05%, 93.64%, and 92.44%, for Models 1, 2, and 3, respectively. These values are close indicating the accuracy of all three models.

As described earlier, we calculated recall and precision for our three models to ensure that the overbalanced training data did not affect the results and also as another means of validating our results. Average values for the three models are presented in Table 4. where even the lowest value of 87.33% for the average precision value for Model 3 is still very good. All other results are above 90% and indicate both that the overbalanced training data did not affect the machine learning results and that the results for all three models are very good.

The next step was using our three designed classifiers on the whole proteome of *L. pneumophila* strain Philadelphia-l to predict effector proteins with results presented in Table 5.

**Table 5.** Comparison of results for the three effector prediction models for *L. pneumophila* strain Philadelphia-1

|  | Number of predicted effector proteins | Number of correctly predicted known: |  | Number of effectors predicted by our models among results for: |  |
| --- | --- | --- | --- | --- | --- |
|  |  | Effectors (316) | Non-effectors (526) | S4TE (302) | Burstein et al. (126) |
| **Model 1** | 760 | 315 (99.7%) | 514 (97.7%) | 273 (90.4%) | 101 (80.2%) |
| **Model 2** | 717 | 300 (94.9%) | 518 (98.5%) | 253 (83.8%) | 100 (79.4%) |
| **Model 3** | 568 | 306 (96.8%) | 521 (99.0%) | 258 (85.4%) | 97 (77.0%) |

The number of predicted effectors is shown in the second column of Table 5. The greatest number of effectors is 760 predicted by Model 1 followed closely by 717 predicted by Model 2. Model 3 predicts 568, considerably fewer and to our knowledge, effector predictions for the three models are greater in number than any previous study for *L. pneumophila* strain Philadelphia-1. As another test of the accuracy of our models, we considered the validated effectors and non-effectors for *L. pneumophila* strain Philadelphia-1 to see which of them were predicted correctly from the test set. These results are shown in the third and fourth columns of Table 5. The lowest of the six results is 94.9% again indicating the overall accuracy of the three models. Model 1 predicts 315 of the 316 validated effector proteins correctly for an accuracy of 99.7%, and Model 3 predicts 521 of 526 non-effector proteins correctly for an accuracy of 99.0%.

We compared our results to effector candidates predicted in two previous studies [5, 6]that focused on *L. pneumophila* strain Philadelphia-1. The first by Burstein et al. experimentally validated 40 new effector proteins and also proposed 126 effector candidates. The second by Meyer et al. proposed 311 candidate effector proteins. These two sets of predicted results shared 45 protein sequences in common, which is 36% of the predicted sequences in [5] and 14% of the predicted sequences in [6]. Our three model comparisons are shown in the fifth and sixth columns of Table 5. Model 1 shares 101 of 126 or 80.2% in common with [5] and 273 of 302 or 90.4% in common with [6] (after removing known non-effectors from their candidates). Interestingly, as shown in Fig. 2, Model 1 also predicted all 45 protein sequences shared by [5] and [6] and also predicted all the 40 new validated effector proteins by [5].

While all three models give good results, the overall results presented in this section indicate that Model 1 is the strongest of the three models. The accuracy metric is the highest, but in addition three of the fold values are above 95%. Recall and precision are most consistent, and comparison with results from previous studies is strongest. The 760 candidate effector proteins for *L. pneumophila* are listed in [S2 Table]. They are also listed in three groups based on the results of the other two models. If predicted by all three models, they are listed in Group 1, by two models in Group 2, and by Model 1 only in Group 3. We assume the first group of 474 has the greatest likelihood of being an effector, the second group of 172 the next most likelihood, and the third group of 114 the next most.

A Venn diagram of the number of candidate effector proteins predicted by Model 1, by Burstein et al. [5], and by Meyer et al. [6] is shown in Fig. 2.

**Figure 2.**
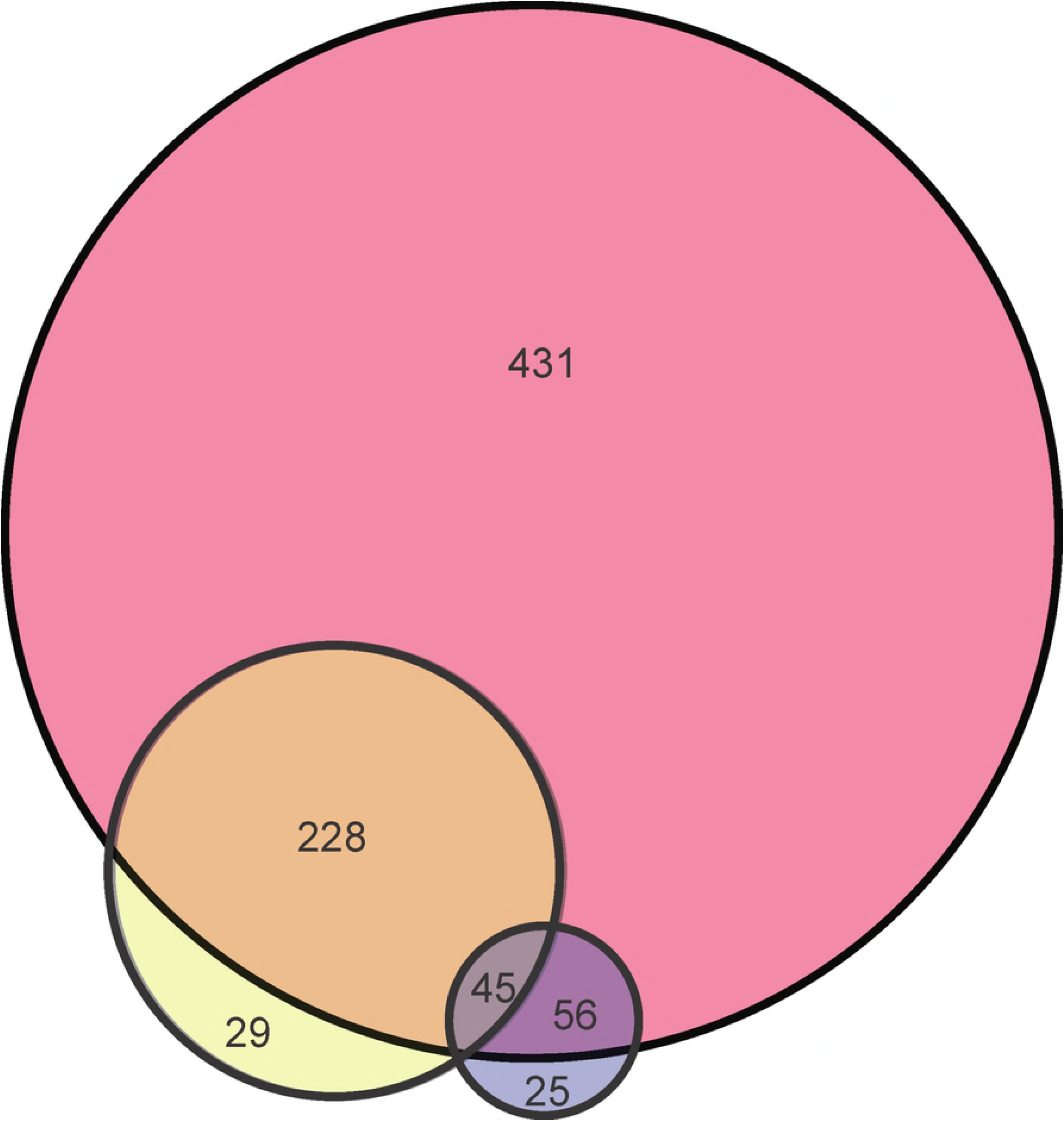
Venn diagram comparing predicted effector proteins for three methods. The pink circle shows the results for Model 1, the yellow circle for the S4TE method, and the blue circle for the method by Burstein et al.

Given the differences shown in this diagram, we conclude that the features used in machine learning predicators are of major importance. More specifically, the reason we predicted more effectors and have more consistent results with previous works, is related to the set of optimal features that we used. This feature set was based on a thorough study of features for the problem of T4SS effector prediction [11, 12]. As the two previous studies developed their models based on a subset of the optimal features, it is likely that they were not able to capture as many effectors. They also had fewer validated effector proteins with which to work.

## Conclusion

In this study, we designed three machine learning classifiers using an optimal set of features and used these classifiers to obtain de novo predictions for effector proteins for *L. pneumophila* strain Philadelphia-1. While all three models were accurate, we found that the strongest model was a straightforward classifier that used all 370 features with a support vector machine. The accuracy, recall, and precision for this model validation, were all greater than 90%. The results of this model compared well with those obtained from two previous research studies predicting more than 80% of the same candidate effector proteins that they did. However, while these older models predicted 126 and 311 candidate effector proteins, our model predicted 760 effector proteins, more than any other model to date, 315 of which have been validated. The reason for this increase in the number of predictions and consistency with previous predictions, is due to the optimal set of features used.

## Supporting information

**S1 File:** Training set composed of known effectors and non-effectors for *L. pneumophila, C. burnettii, Brucella* spp., and *Bartonella* spp.

**S2 File:** Test set composed of all protein sequences from the whole proteome for *L. pneumophila* strain Philadelphia-1.

**S1 Table:** The 370 features used in the three machine learning models developed for this study.

**S2 Table:** The set of 760 de novo effector proteins predicted by Model 1 for *L. pneumophila* strain Philadelphia-1.

## References

1. Han N, Yu W, Qiang Y, Zhang W. T4SP Database 2.0: An Improved Database for Type IV Secretion Systems in Bacterial Genomes with New Online Analysis Tools. Computational and Mathematical Methods in Medicine. 2016; 2016, 9415459. (10.1155/2016/9415459)

2. Voth DE, Broederdorf, LJ, Graham JG. Bacterial Type IV Secretion Systems: Versatile Virulence Machines. Future Microbiology. 2012; 7(2), 241–257. (10.2217/fmb.11.150)

3. Voth DE, Beare PA, Howe D, Sharma UM, Samoilis G, Cockrell DC, Omsland A, Heinzen RA. The Coxiella burnetii Cryptic Plasmid Is Enriched in Genes Encoding Type IV Secretion System Substrate. Journal of Bacteriology. 2010; 193(7), 1493–1503. (doi: 10.1128/JB.01359-10)

4. Abby SS, Cury J, Guglielmini J, Néron B, Touchon M, Rocha EPC. Identification of protein secretion systems in bacterial genomes. Scientific Reports. 2016; 6. (doi: 10.1038/srep23080).

5. Burstein D, Zusman T, Degtyar E, Viner R, Segal G, Pupko T. Genome-Scale Identification of Legionella pneumophila Effectors Using a Machine Learning Approach. The International Journal of Biochemistry and Cell Biology. 2009; 5(7). (https://doi.org/10.1371/journal.ppat.1000508)

6. Meyer D, Noroy C, Moumene A, Raffaele S, Albina E, Vachiery N. Searching algorithm for type IV secretion system effectors 1.0: a tool for predicting type IV effectors and exploring their genomic context. Nucleic Acids Research. 2013; 41(20), 9218–9229. (doi: 10.1093/nar/gkt718)

7. Zou L, Nan C, Hu F. 2013. Accurate prediction of bacterial type IV secreted effectors using amino acid composition and PSSM profiles. Bioinformatics 29(24), 3135–3142. (doi: 10.1093/bioinformatics/btt554)

8. Yu L, Guo Y, Li Y, Li G, Li M, Luo J, Xiong W, Qin W. 2013. SecretP: identifying bacterial secreted proteins by fusing new features into Chous pseudo-amino acid composition. J Theor Biol. 267, 1–6. (doi: 10.1016/j.jtbi.2010.08.001)

9. Wang Y, Wei X, Bao H, Liu S. Prediction of bacterial type IV secreted effectors by C-terminal features. BMC Genomics 2014; 15(50). (doi: 10.1186/1471-2164-15-50)

10. Lockwood S, Voth D, Brayton K, Beare P, Brown W, Heinzen R, Broschat S. Identification of Anaplasma marginale Type IV Secretion System Effector Proteins. PLoS ONE. 2011; 6(11), e27724. (https://doi.org/10.1371/journal.pone.0027724)

11. Esna Ashari Z, Brayton K, Broschat S. Determining Optimal Features for Predicting Type IV Secretion System Effector Proteins for Coxiella burnetii. Proceedings of 8th ACM BCB conference. 2017; 346–351.

12. Esna Ashari Z, Dasgupta N, Brayton K, Broschat S. An optimal set of features for predicting type IV secretion system effector proteins for a subset of species based on a multi-level feature selection approach. PLoS ONE 2018; 13, e0197041. (https://doi.org/10.1371/iournal.pone.0197041)

13. Bruggemann H, Cazalet C, Buchrieser C. Adaptation of Legionella pneumophila to the host environment: role of protein secretion, effectors and eukaryotic-like proteins. Current Opinion in Microbiology. 2006; 9(1), 86–94.

14. Cazalet C, Rusniok R, Bruggemann H, Zidane N, Magnier A, Ma L, Tichit M, Jarraud S, Bouchier C, Vandenesch F, Kunst F, Etienne J, Glaser P, Buchrieser C. Evidence in the Legionella pneumophila genome for exploitation of host cell functions and high genome plasticity. Nature Genetics. 2004; 36(11), 1165–1173.

15. Chen J, Suwwan de Felipe K, Clarke M, Lu H, Anderson O, Segal G, Shuman H. Legionella Effectors That Promote Nonlytic Release from Protozoa. Science. 2004; 303(5662), 1358–1361. (doi: 10.1126/science.1094226)

16. Suwwan de Felipe K, Pampou S, Jovanovic O, Pericone C, Ye S, Kalachikov S, Shuman H. Evidence for Acquisition of Legionella Type IV Secretion Substrates via Interdomain Horizontal Gene Transfer. Journal of Bacteriology. 2005; 187(22), 7716–7726.

17. Conover G, Derre I, Vogel J, RR I. The Legionella pneumophila LidA protein: a translocated substrate of the Dot/Icm system associated with maintenance of bacterial integrity. Molecular Microbiology. 2003; 48(2), 305–321.

18. Laguna R, Creasey E, Li Z, Valtz N, Isberg R. A Legionella pneumophila-translocated substrate that is required for growth within macrophages and protection from host cell death. Proceedings of the National Academy of Sciences. 2006; 103(49), 18745–18750.

19. Bardill J, Miller J, Vogel J. IcmS-dependent translocation of SdeA into macrophages by the Legionella pneumophila type IV secretion system. Molecular Microbiology. 2005; 56(1), 90–103.

20. Ninio S, Zuckman-Cholon D, Cambronne E, Roy C. The Legionella IcmS-IcmW protein complex is important for Dot/Icm-mediated protein translocation. Molecular Microbiology. 2005; 55(3), 912–926.

21. Altman E, Segal G. The Response Regulator CpxR Directly Regulates Expression of Several Legionella pneumophila icm/dot Components as Well as New Translocated Substrates. Future Microbiology. 2008; 190(6), 1985–1996. (doi:10.1128/JB.01493-07)

22. Zusman T, Aloni G, Halperin E, Kotzer H, Degtyar E, Feldman M, Segal G. The response regulator PmrA is a major regulator of the icm/dot type IV secretion system in Legionella pneumophila and Coxiella burnetii. Molecular Microbiology. 2007; 63(5), 1508–1523.

23. Zusman T, Degtyar E, Segal G. Identification of a Hypervariable Region Containing New Legionella pneumophila Icm/Dot Translocated Substrates by Using the Conserved icmQ Regulatory Signature. Infection and Immunity. 2008; 76(10), 4581–4591. (doi:10.1128/IAI.00337-08)

24. Suwwan de Felipe K, Glover R, Charpentier X, Anderson O, Reyes M, Pericone C, Shuman H. Legionella Eukaryotic-Like Type IV Substrates Interfere with Organelle Trafficking. PLoS Pathogens. 2008; 4(8). (doi: 10.1371/journal.ppat.1000117)

25. Heidtman M, Chen E, Moy M, Isberg R. Large scale identification of Legionella pneumophila Dot/Icm substrates that modulate host cell vesicle trafficking pathways. Cellular Microbiology. 2009; 11(2), 230–248. (doi:10.1111/j.1462-5822.2008.01249.x)

26. Shohdy N, Efe J, Emr S, Shuman H. Pathogen effector protein screening in yeast identifies Legionella factors that interfere with membrane trafficking. Proceedings of the National Academy of Sciences. 2005; 102(13).

27. Nagai H, Cambronne E, Kagan J, Amor J, Kahn R, Roy C. A C-terminal translocation signal required for Dot/Icm-dependent delivery of the Legionella RalF protein to host cells. Proceedings of the National Academy of Sciences. 2005; 102(3), 826–831.

28. Stormo G, Schneider T, Gold L, Ehrenfeucht A. Use of the ‘Perceptron’ algorithm to distinguish translational initiation sites in E. cosali. Nucleic Acids Research. 1982; 10(9), 2997–3011.

29. Cortes C, Vapnik V. Support-vector networks. Machine Learning 1995; 20(3), 273–297.

30. Crammer K, Singer Y. On the Algorithmic Implementation of Multiclass Kernel-based Vector Machines. JMLR. 2001; 2, 265–292.

31. Perry J W, Kent A, Berry M. Machine literature searching X. Machine language; factors underlying its design and development. American Documentation. 1955; 6(4), 242–254.

